# Retroelement Hypomethylation Links Hypoxia Signaling, Immune Phenotypes, and Survival in Clear Cell Renal Cell Carcinoma

**DOI:** 10.64898/2026.05.01.722263

**Authors:** Chinaza F. Nnam, Erick A. Mboya, Yiping Li, Minghui Zhang, Fred Kolling, Laurent Perrard, Thomas J. Palys, Elizabeth Pflugradt, Patricia A. Pioli, Marc Ernstoff, John D. Seigne, Jason R. Pettus, Bing Ren, Li Song, Brock C. Christensen, Lucas A. Salas

**Affiliations:** Department of Cancer Biology, Guarini School of Graduate and Advanced Studies, Dartmouth College, Hanover, NH, USA; Department of Epidemiology, Geisel School of Medicine, Dartmouth College, Lebanon, NH, USA; Department of Microbiology and Immunology, Guarini School of Graduate and Advanced Studies, Dartmouth College, Hanover, NH, USA; Department of Biomedical Data Science, Geisel School of Medicine, Dartmouth College, Lebanon, NH, USA; Department of Molecular and Systems Biology, Geisel School of Medicine, Dartmouth College, Lebanon, NH, USA; Dartmouth Hitchcock Medical Center. Lebanon, New Hampshire; Dartmouth Cancer Center Shared Resources

**Author notes:** Corresponding: Lucas A. Salas.

**Keywords:** clear cell renal cell carcinoma, DNA methylation, retroelements, prognosis, immune response, hypoxia signaling

## Abstract

**Background:** Retrotransposable elements (RE) comprise approximately 45% of the human genome and are typically repressed by DNA methylation to preserve genomic integrity. In cancer, global DNA hypomethylation can lead to RE derepression, resulting in genomic instability and activation of innate immune pathways through viral mimicry. While individual RE classes have been examined in clear cell renal cell carcinoma (ccRCC), the integrated epigenetic landscape of multiple RE families and their clinical relevance remain incompletely characterized.

**Methods:** We performed a genome-wide prediction of DNA methylation across three major RE classes (Alu, LINE-1, and LTR elements) using a validated computational framework applied to Illumina methylation array data from two independent ccRCC tumor cohorts. Integrated unsupervised clustering of RE methylation profiles was used to define the epigenetic subtypes. Associations with clinicopathologic variables, tumor immune microenvironment composition (DNA Methylation-derived), hypoxia signaling, innate immune activation, and overall survival were evaluated. Prognostic relevance was assessed using multivariable Cox regression models adjusting for age, sex, AJCC stage or AUA risk group, and immune and angiogenic tumor microenvironment features. Key findings were then externally validated in CPTAC-ccRCC and independently replicated in an institutional Dartmouth Cancer Center (DCC) cohort with matched methylation and RNA-sequencing data.

**Results:** Integrated clustering identified three reproducible RE methylation subtypes, Repressed, Transient, and Active. In the discovery cohort, the Active subtype showed significantly worse overall survival than the Repressed subtype, with a graded survival pattern across RE methylation states that persisted after multivariable adjustment. RE hypomethylation was associated with reduced EPAS1 (HIF2A) expression, increased immune infiltration, elevated PD-1 expression, and heightened cGAS-STING and interferon signaling, consistent with an immune-inflamed yet immunosuppressed tumor state. In the external CPTAC validation cohort, RE methylation subtypes recapitulated key molecular features and showed supportive survival trends. In the independent DCC replication cohort, an Active RE state was again associated with poorer survival, lower EPAS1 expression, increased PD-1 expression, greater CD8 T-cell and Treg infiltration, and elevated T-cell exhaustion signatures, supporting the reproducibility of the prognostic and immune-exhausted phenotype across cohorts.

**Conclusions:** We identified RE methylation subtypes with distinct molecular, immunologic, and prognostic features in ccRCC. External validation in CPTAC and independent replication in DCC support the robustness of this RE methylation framework across large-scale and institutional cohorts. These findings highlight the prognostic potential of RE methylation profiles and support their integration into molecular classification strategies to improve risk stratification in ccRCC.

## 1. Introduction

An estimated 45% of the human genome consists of DNA sequences derived from transposable elements, which are remnants of ancient mobile genetic elements that have integrated throughout the genome^1–3^. Retrotransposons (RE) are the most abundant class of transposable elements and replicate through an RNA intermediate. Based on structural features, RE are broadly categorized into two groups: long terminal repeat (LTR) elements and non-LTR elements^4,5^. The LTR group includes human endogenous retroviruses (HERVs), which comprise about 8% of the genome^6^. Although most HERVs are now immobile due to accumulated mutations and internal deletions, their long terminal repeat regions retain the potential to influence gene expression through promoter or enhancer activity^7^. Non-LTR elements include Long and Short Interspersed Nuclear Elements (LINEs and SINEs, respectively). LINE-1 (L1) elements, which make up nearly 17% of the genome, remain capable of autonomous retrotransposition through the activity of encoded reverse transcriptase and endonuclease proteins^8^. In contrast, Alu elements, the most abundant SINEs, are non-autonomous and rely on the enzymatic machinery of L1 elements for mobilization^3,4,6,10^.

In normal somatic cells, RE activity is tightly controlled by DNA methylation and other epigenetic mechanisms that maintain genomic integrity^10,11^. However, aberrant global hypomethylation, as occurs in carcinogenesis, can lead to RE reactivation, resulting in genomic instability and chromatin remodeling^5,12^. Such reactivation can also engage innate and adaptive immune pathways, thereby influencing tumor–immune interactions^13^. One mechanism linking RE activation to immune signaling is viral mimicry, a cellular state in which endogenous nucleic acids trigger antiviral responses typically reserved for exogenous viral infections^6,14^. Among other mechanisms, HERV activation can induce cytosolic nucleic acid accumulation, which stimulates immune signaling pathways such as cGAS–STING^15^. In addition, L1 retroelements have emerged as potent inducers of the cGAS–STING axis, a pathway that has demonstrated prognostic relevance in several cancers, including clear cell renal cell carcinoma (ccRCC)^16^.

ccRCC, the most common histologic subtype of kidney cancer, is characterized by alterations in the von Hippel–Lindau (*VHL*) tumor suppressor gene that result in pseudohypoxia and transcriptional activation of hypoxia-inducible factors (HIFs)^17^. This pseudohypoxic state drives metabolic reprogramming, immune evasion, and extensive epigenetic remodeling, including widespread changes in DNA methylation^18,16^. Recent evidence suggests that hypoxia and metabolic stress can promote transposable element derepression, potentially through impaired maintenance of methylation and increased oxidative stress^20^. In ccRCC, RE activation and immune pathway dysregulation are particularly relevant, as ccRCC is characterized by a highly infiltrated tumor microenvironment with active immune checkpoint signaling^21^. Studies have linked RE reactivation with immune dysregulation in ccRCC. HERV-H LTR-associating protein 2 (HHLA2), which has been shown to inhibit T-cell receptor signaling, is significantly elevated in ccRCC^22^. In addition, a subset of HERV elements (referred to as group 2 HERV) associated with enhanced immune suppression has been linked to downregulation of the RIG-I–like receptor pathway^23^. Higher cGAS–STING pathway activity, which may in part be attributed to HERV activation, has also been associated with worse prognosis in ccRCC^24^. These observations suggest that RE activation may shape tumor immune phenotypes and influence clinical outcomes in ccRCC. However, the landscape of RE methylation in ccRCC and its relationship with tumor biology and patient survival remains poorly understood.

Most prior studies have focused on a single RE class, often assessed using locus-nonspecific assays such as pyrosequencing, which cannot distinguish between individual RE copies or genomic contexts^25,26^. In contrast, recent computational advances now enable high-resolution, genome-wide estimation of methylation across specific RE families, subfamilies, and genomic locations, allowing integrative analysis of multiple RE types within the same tumor. Because maintenance of DNA methylation is essential for retrotransposon repression, profiling methylation across different RE classes can provide critical insights into their epigenetic regulation and potential reactivation in cancer^5,27,28^. In this study, our goal was to perform a comprehensive analysis of methylation across three major RE classes (LTRs, L1s, and Alus) in ccRCC. Using multi-omics clustering techniques, we aimed to identify RE-defined tumor subgroups in the TCGA-KIRC cohort and evaluate their associations with clinicopathologic features, tumor immune composition, and patient survival. Given the extensive epigenetic and metabolic reprogramming driven by VHL loss and hypoxia in ccRCC, we hypothesize that RE methylation patterns capture key aspects of tumor biology and may serve as independent prognostic indicators beyond established clinical and molecular factors. Importantly, we further sought to validate the robustness and generalizability of these RE-defined subtypes in an external cohort, CPTAC-ccRCC, and to independently replicate the key findings in an institutional Dartmouth Cancer Center cohort from the Dartmouth Renal Cancer Tumor Bank, thereby establishing their reproducibility across platforms and clinical contexts.

## 2. Methods

### 2.1. Datasets

DNA methylation data for primary clear cell renal cell carcinoma (ccRCC) tumors were obtained from two independent cohorts: The Cancer Genome Atlas Kidney Renal Clear Cell Carcinoma (TCGA-KIRC) and the Clinical Proteomic Tumor Analysis Consortium (CPTAC)^26,30^. Raw IDAT files generated using the Illumina HumanMethylation450K (TCGA-KIRC) were downloaded through the TCGAbiolinks R package^31^ together with corresponding demographic and clinical annotations. For external validation, raw HumanMethylationEPIC v1 array data files from the CPTAC-ccRCC cohort were also downloaded as in the TCGA-KIRC cohort. For transcriptomic analyses, gene expression data in transcripts per million (TPM) were retrieved for both cohorts and log2-transformed. For independent validation, we leveraged an ongoing institutional multiomics ccRCC cohort at Dartmouth Cancer Center (DCC), including 105 tumors with matched raw HumanMethylationEPIC v1 array data and raw paired-end RNA-sequencing reads. DCC samples were collected between 1994 and 2014 under institutional review board approval (IRB# STUDY02000672). TCGA-KIRC somatic mutation and copy number alteration data for key ccRCC driver genes (*VHL, BAP1, SETD2,* and *PBRM1*) were obtained from cBioPortal^32^.

### 2.2 DCC Cohort Library Preparation, Sequencing, and Methylation Profiling

Total RNA was extracted from frozen tissue using Qiagen RNeasy kits according to the manufacturer’s instructions. RNA integrity was assessed using an Agilent 2100 Bioanalyzer, and only samples with RNA integrity number (RIN) ≥ 7 were advanced to library preparation. Ribo-depleted libraries were generated using the Illumina TruSeq Stranded mRNA kit (ribo-depletion workflow) and sequenced on an Illumina NovaSeq 6000 to generate 2 × 150 bp paired-end reads, targeting ∼50 million paired-end reads per sample.

### 2.3. DCC Cohort RNA-seq Preprocessing and Quantification

All raw sequencing reads were processed using a standardized Nextflow^33^ pipeline (v24.10.5). Adapter sequences and low-quality bases were removed using TrimGalore!, and read quality metrics were summarized using FastQC and MultiQC. Reads were aligned to the human reference genome GRCh38.p13 using STAR (v2.7)^34^. Gene-level counts were quantified from aligned BAM files using featureCounts. Transcript-level quantification was performed using Salmon (v1.5)^35^ in mapping-based mode against the Ensembl v113 transcriptome to generate transcript-level estimated counts and TPM values. For downstream analyses requiring normalized expression, TPM values were log2-transformed as log2(TPM + 1). Gene-level TPM values were derived from Salmon outputs when needed to facilitate consistent expression scaling across cohorts. To identify potential sample contamination, Kraken2 and Bracken were used to screen reads for non-human content. Samples exceeding 1% bacterial, viral, or fungal reads were excluded prior to downstream statistical analyses.

### 2.4 DNA Methylation Preprocessing and Cell Type Deconvolution

For all datasets, raw IDAT files were preprocessed using the ENmix pipeline (R v4.5.1)^36^. Specifically, we applied a stringent pOOBAH detection p-value threshold of 0.05, required at least three beads per probe, and removed probes with fluorescence signal intensities more than three standard deviations from the mean of the bisulfite conversion control probes. Probes failing these criteria were masked as missing. Probes with more than 5% missing values across samples were excluded, and no imputation was performed at this stage to avoid introducing artificial methylation variability. Samples with more than 10% missing CpG values or with beta value distributions deviating by more than three times the interquartile range from the cohort median were excluded as these could represent technically biased or unsuccessful assays. CpG loci known to cross-react with multiple genomic loci or overlap polymorphic sites with minor allele frequency greater than 1% in either global or African populations were masked to minimize SNP interference in methylation quantification^37^. All non-CpG (CpH) probes were removed due to low biological variability^38^, and probes mapping to the X chromosome were excluded in this initial pass to avoid sex-linked methylation bias related to dosage compensation and imprinting effects^36^. For CPTAC but not the replication cohort DCC, beta values, EPIC probes were restricted to those overlapping with the 450K array. After filtering, high-quality CpG probes for CPTAC-ccRCC were retained for downstream analyses.

To ensure all samples represented ccRCC, tumor histologic identity was confirmed using the Hierarchical Tumor Artificial Intelligence Classifier (HiTAiC), a deep learning based classifier^40^. Samples that were mismatched were excluded. The hierarchical tumor immune microenvironment epigenetic deconvolution framework (HiTIMED) was used to estimate tumor microenvironment proportions of 17 cell types in both datasets^41^. A total of 310 TCGA-KIRC, 275 CPTAC, and 116 DCC tumor samples remained after quality control and HiTAIC confirmation.

### 2.5 Prediction of Alu, L1, and LTR Methylation in ccRCC

DNA methylation levels of the three major classes of REs, LTR, LINE-1 (L1), and Alu, were predicted using a Random Forest-based algorithm implemented in the REMP (Repetitive Element Methylation Prediction) R package^42^. Preprocessed beta values were first processed using the grooMethy() function to identify and correct data irregularities, including missing, zero, or infinite values. For each RE class, an annotated database was generated using the initREMP() function. The hg38 genome build was specified for alignment, and annotation databases were downloaded from the UCSC website using annotation.option.source = “AH” for the Illumina HumanMethylation450K TCGA-KIRC cohort and “UCSC” for the Illumina HumanMethylationEPIC CPTAC-ccRCC cohort.

RE methylation prediction was performed using the remp() function with a 1,000 bp flanking window. Predicted values with quality scores below 1.7 or missing rates greater than 20% were excluded according to REMP package recommendations. CpG-level methylation predictions were aggregated using the rempAggregate() function, requiring at least two valid CpG estimates per RE element. Predicted methylation density distributions were visually inspected for outliers. For TCGA, two samples were removed, and the prediction pipeline was rerun. Finally, predicted RE loci were annotated using rempAnnot(), resulting in genome-wide methylation estimates for LTR, L1, and Alu retroelements.

### 2.6 Integrated RE Clustering

Following RE methylation prediction and aggregation, missing values were imputed using the k-nearest neighbors algorithm implemented in the impute.knn() function from the impute package^43^. Each missing value was estimated from the methylation profiles of the 15 most similar probes based on Euclidean distance. Feature selection was performed using the median absolute deviation to identify the 1,000 most variable LTR, L1, and Alu retroelements for clustering. To identify molecular subgroups defined by RE methylation patterns, multi-omics clustering was conducted using six integrative approaches implemented in the MOVICS framework^44^: Neighborhood-based Multi-Omics Clustering (NEMO), Similarity Network Fusion (SNF), Integrated Non-Negative Matrix Factorization (IntNMF), moCluster, Consensus Clustering, and Cluster-of-Clusters Analysis (COCA)^45–50^. NEMO and SNF construct sample similarity networks across RE methylation layers. NEMO integrates concordant local neighborhood relationships and accommodates partially missing data, whereas SNF iteratively diffuses similarity across layer-specific networks to generate a fused similarity network for clustering. Cluster-of-Clusters Analysis integrates independently derived cluster assignments into a meta-clustering framework. Integrated Non-Negative Matrix Factorization and moCluster perform joint matrix factorization to extract latent components modeling shared and layer-specific variation across RE classes. Consensus Clustering performs repeated subsampling of combined feature matrices derived from individual RE layers. The optimal number of clusters was determined using the Cluster Prediction Index (CPI) across candidate k values (k= 2:8). Heatmaps of the 1,000 most variable RE positions per class were generated from M values using the getMoHeatmap() function in MOVICS and the ComplexHeatmap package and were annotated with cluster assignments. Heatmaps based on hierarchical clustering were generated using the pheatmap package.

### 2.7 Immune-Related Gene Signature Scoring

Gene sets representing cGAS–STING pathway activation, interferon-stimulated genes, and T-cell progenitor and terminal exhaustion states were curated from previously published studies and reviews (Table S1)^24,51,52^. For each tumor sample, enrichment scores were computed using single-sample Gene Set Enrichment Analysis implemented in the Gene Set Variation Analysis package^53^. Scoring was performed using default parameters.

### 2.8 Statistical and Survival Analysis

Samples lacking complete clinical information were excluded from survival analyses. The final survival dataset included 308 TCGA-KIRC, 243 CPTAC-ccRCC (n = 167 unique individuals) and 116 DCC tumor samples. For TCGA-KIRC, overall survival was defined as days to death or last follow-up. Associations between RE-defined subtypes and clinical covariates were evaluated using one-way analysis of variance for continuous variables and chi-square tests for categorical variables. Associations between RE-defined subgroups and overall survival were evaluated using Cox proportional hazards regression. Models were adjusted sequentially for age, sex, AJCC stage, immune cell composition, angiogenic features, and AUA risk stratification. Proportional hazards assumptions were assessed using Schoenfeld residuals. For CPTAC, Cox models incorporated a robust sandwich variance estimator to account for duplicate tumor samples from the same individuals. Differential methylation analyses were performed with and without adjustment for clinical and tumor microenvironment covariates. Survival analyses were conducted using the survival package in R^54^. Associations between RE clusters and molecular phenotypes, including *HIF1A* and *EPAS1* mRNA expression, cGAS–STING activation scores, and interferon-stimulated gene signatures, were tested using Wilcoxon rank-sum tests. All analyses and visualizations were conducted in R (v4.5.1).

## 3. Results

### 3.1. Study Populations

Two independent clear cell renal cell carcinoma (ccRCC) cohorts were analyzed: TCGA-KIRC (n = 308, discovery) and CPTAC-ccRCC (n = 167, validation). In TCGA-KIRC, the mean age at diagnosis was 61.3 (±11.62) years, with a predominance of male patients (65.9%). The CPTAC cohort showed similar demographic patterns, with a mean age of 60.8 (±11.8) years, and a comparable male predominance (71%). Only 16.3% of CPTAC patients had died at last follow-up, reflecting fewer events and generally favorable outcomes relative to TCGA (33%) (Table 1).

**Table 1.** Clinical Characteristics of TCGA-KIRC, CPTAC-CCRCC and DCC cohorts.

| Covariates | Groups | TCGA-KIRC |  | p | CPTAC-ccRCC |  | p | DCC |  | p |
| --- | --- | --- | --- | --- | --- | --- | --- | --- | --- | --- |
|  |  | Alive<br>N=207 | Dead<br>N=101 |  | Alive<br>N=140 | Dead<br>N=27 |  | Alive<br>N=66 | Dead<br>N=49 |  |
| Age(mean (SD)) |  | 59.69<br>(11.27) | 64.61<br>(11.69) | <0.001 | 60.16<br>(11.86) | 64.11<br>(11.09) | 0.111 | 60.23 (10.33) | 63.68 (12.50) | 0.120 |
| Sex, n (%) | Female | 71 (34.3) | 34 (33.7) | 1.000 | 99 (70.7) | 18 (66.7) | 0.849 | 27 (41.5) | 11 (22.4) | 0.052 |
|  | Male | 136 (65.7) | 67 (66.3) |  | 41 (29.3) | 9 (33.3) |  | 38 (58.5) | 38 (77.6) |  |
| Stage, n (%) | I | 132 (63.8) | 19 (18.8) | <0.0001 | 83 (59.3) | 4 (14.8) | <0.001 | 39 (63.9) | 13 (33.3) | <0.001 |
|  | II | 20 (9.7) | 8 (7.9) |  | 13 (9.3) | 3 (11.1) |  | 9 (14.8) | 3 (7.7) |  |
|  | III | 43 (20.8) | 28 (27.7) |  | 36 (25.7) | 14 (51.9) |  | 10 (16.4) | 9 (23.1) |  |
|  | IV | 12 (5.8) | 46 (45.5) |  | 8 (5.7) | 6 (22.2) |  | 3 (4.9) | 14 (35.9) |  |
| Grade, n (%) | G1 | 7 (3.4) | 0 (0.0) | <0.0001 | 13 (9.3) | 2 (7.4) | 0.004 | 5 (7.9) | 0 (0.0) | 0.076 |
|  | G2 | 113 (54.6) | 20 (19.8) |  | 81 (57.9) | 9 (33.3) |  | 31 (49.2) | 14 (35.9) |  |
|  | G3 | 76 (36.7) | 43 (42.6) |  | 40 (28.6) | 10 (37.0) |  | 17 (27.0) | 13 (33.3) |  |
|  | G4 | 11 (5.3) | 38 (37.6) |  | 6 (4.3) | 6 (22.2) |  | 10 (15.9) | 12 (30.8) |  |
| AUA Risk<br>(Binary), n<br>(%) | Intermediate_high |  |  |  |  |  |  | 33 (55.0) | 30 (76.9) | 0.045 |
|  | Low |  |  |  |  |  |  | 27 (45.0) | 9 (23.1) |  |

### 3.2. Integrative clustering of Alu, L1, and LTR methylation identifies three robust retroelement epigenetic subtypes in ccRCC

After filtering, 348,771 CpGs for TCGA-KIRC dataset were used for RE methylation prediction and aggregation. A total of 9,270 RE positions were retained after aggregation, including 6,361 Alu, 1,271 L1, and 1,638 LTR positions. In addition, missing values, representing 6% of total beta values, were imputed. To define the retroelement (RE) centered epigenetic landscape of ccRCC, we benchmarked six multiomics clustering frameworks, NEMO, SNF, moCluster, IntNMF, COCA, and Consensus Clustering, on integrated REMP predicted Alu, L1, and LTR methylation profiles. Gap statistic profiling using the getClustNum() function supported an optimal cluster number of k = 3 (Figure S1). Across the six approaches, only NEMO and SNF consistently generated well-separated, homogeneous clusters, exhibiting low within-cluster dispersion and high between-cluster divergence. While consensus-based approaches (ConsensusClustering, COCA) tended to highlight outlier partitions, IntNMF produced evenly balanced clusters, reflecting its inherent preference for size-balanced solutions rather than structures optimized for underlying biological variation (Figure S2). Apart from NEMO and SNF, the remaining methods showed limited capacity to capture the asymmetric, potentially biologically meaningful variation in RE methylation characteristic of tumor-intrinsic evolutionary pressures and heterogeneity. Because NEMO’s network-fusion architecture preserves inter-layer correlations, it was selected as the primary integrative framework for downstream molecular and clinical analyses.

Application of NEMO to the full RE methylomes in TCGA-KIRC revealed three highly distinct and biologically coherent methylation subtypes (Figure 1). A large Repressed subtype (n = 171) displayed uniformly elevated methylation across Alu, L1, and LTR elements, consistent with widespread silencing of repetitive elements and maintenance of genomic repression. A smaller Active subtype (n = 80) exhibited pervasive hypomethylation across all RE classes, suggestive of global derepression and potential ERV transcriptional activation. An intermediate Transient subtype (n = 57) showed partial demethylation, representing a possible transitional epigenetic state between complete repression and activation (Figure 1). Clinical characteristics differed significantly across the three RE-defined subtypes (Table 2).

**Figure 1.**
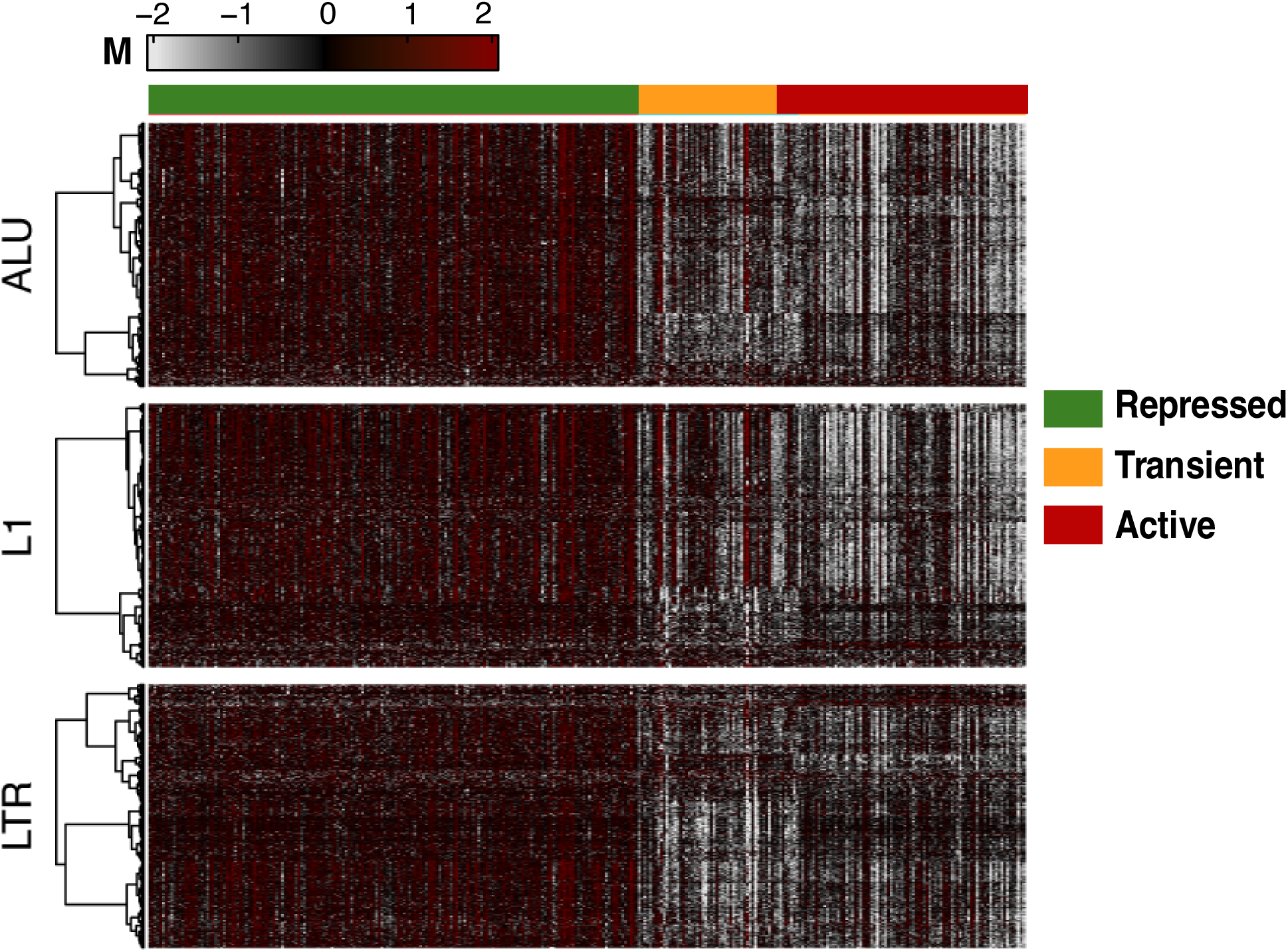
Integrated clustering of retroelement methylation identifies three distinct epigenetic subtypes in ccRCC. Network-fusion–based Neighborhood-based Multi-Omics Clustering (NEMO) applied to integrated REMP-predicted Alu, LINE-1 (L1), and LTR methylation profiles stratified TCGA-KIRC tumors into three distinct and internally homogeneous subgroups. The Repressed subtype is characterized by globally high retroelement methylation, the Active subtype exhibits widespread hypomethylation consistent with retroelement derepression, and the Transient subtype displays intermediate methylation levels.

**Table 2.** Distribution of clinical characteristics across RE-defined subtype. The Repressed group is enriched for early-stage and lower-grade tumors, whereas the Transient and Active subtypes show higher proportions of advanced-stage disease and high-grade tumors. Age differed significantly across subtypes (one-way ANOVA, *p* = 0.009), while sex distribution did not differ (chi-square test). Tumor stage and grade were strongly stratified across RE subtypes (chi-

| Groups |  | Repressed<br>n=171 | Transient<br>n=57 | Active<br>n=80 | p |
| --- | --- | --- | --- | --- | --- |
| Age, mean<br>(SD) (yrs) |  | 59.58 (12.08) | 64.49 (11.27) | 62.73 (10.25) | 0.009 |
| Sex n (%) | Female | 54 (31.6) | 22 (38.6) | 29 (36.2) | 0.56 |
|  | Male | 117 (68.4) | 35 (61.4) | 51 (63.7) |  |
| Stage, n (%) | I | 116 (67.8) | 19 (33.3) | 16 (20.0) | <0.001 |
|  | II | 14 (8.2) | 6 (10.5) | 8 (10.0) |  |
|  | III | 26 (15.2) | 14 (24.6) | 31 (38.8) |  |
|  | IV | 15 (8.8) | 18 (31.6) | 25 (31.2) |  |
| Grade, n (%) | G1 | 6 (3.5) | 1 (1.8) | 0 (0.0) | <0.001 |
|  | G2 | 94 (55.0) | 22 (38.6) | 17 (21.2) |  |
|  | G3 | 59 (34.5) | 24 (42.1) | 36 (45.0) |  |
|  | G4 | 12 (7.0) | 10 (17.5) | 27 (33.8) |  |

square tests, both $p < 0.001$ .

Patients in the Transient and Active groups were slightly older on average than those in the Repressed group (mean ages 64.5 and 62.7 vs. 59.6 years; *p* = 0.009). Sex distribution did not differ across subtypes (*p* = 0.560). By contrast, tumor stage and grade showed marked stratification. Early-stage disease (Stage I) was enriched in the Repressed group (67.8%) but substantially less common in the Transient (33.3%) and Active (20.0%) groups, whereas advanced tumors (Stage III– IV) were more frequent in the Transient (56.2%) and Active (70.0%) subtypes (*p* < 0.001). Tumor grade followed a similar pattern: lower-grade tumors (G1–G2) predominated in the Repressed subtype (58.5%), while high-grade tumors (G3–G4) were increasingly represented in the Transient (59.6%) and especially the Active group (78.8%; *p* < 0.001). The concordant subtype structure across three evolutionarily distinct RE families, coupled with clear clinical stratification, underscores that NEMO captures a shared genome-wide derepression axis with meaningful phenotypic consequences rather than class-specific noise or technical variability.

### 3.3. Active RE methylation subtype is associated with worse survival independent of clinical stage, AUA risk group, and TME composition

We further assessed whether RE methylation subtypes stratify patient survival. A Kaplan–Meier analysis showed clear and significant survival differences among the three NEMO-defined subtypes (Figure 2). Patients in the Repressed subtype experienced the most favorable outcomes and did not reach 50% mortality during follow-up. The Transient subtype showed intermediate survival with a median overall survival of 7.02 years (95% CI: 4.5–not reached), whereas the Active subtype had the poorest prognosis, with a median overall survival of 3.7 years (95% CI: 2.4–6.1). Survival differences were highly significant (log-rank p < 0.0001), demonstrating a graded risk pattern of Repressed > Transient > Active.

**Figure 2.**
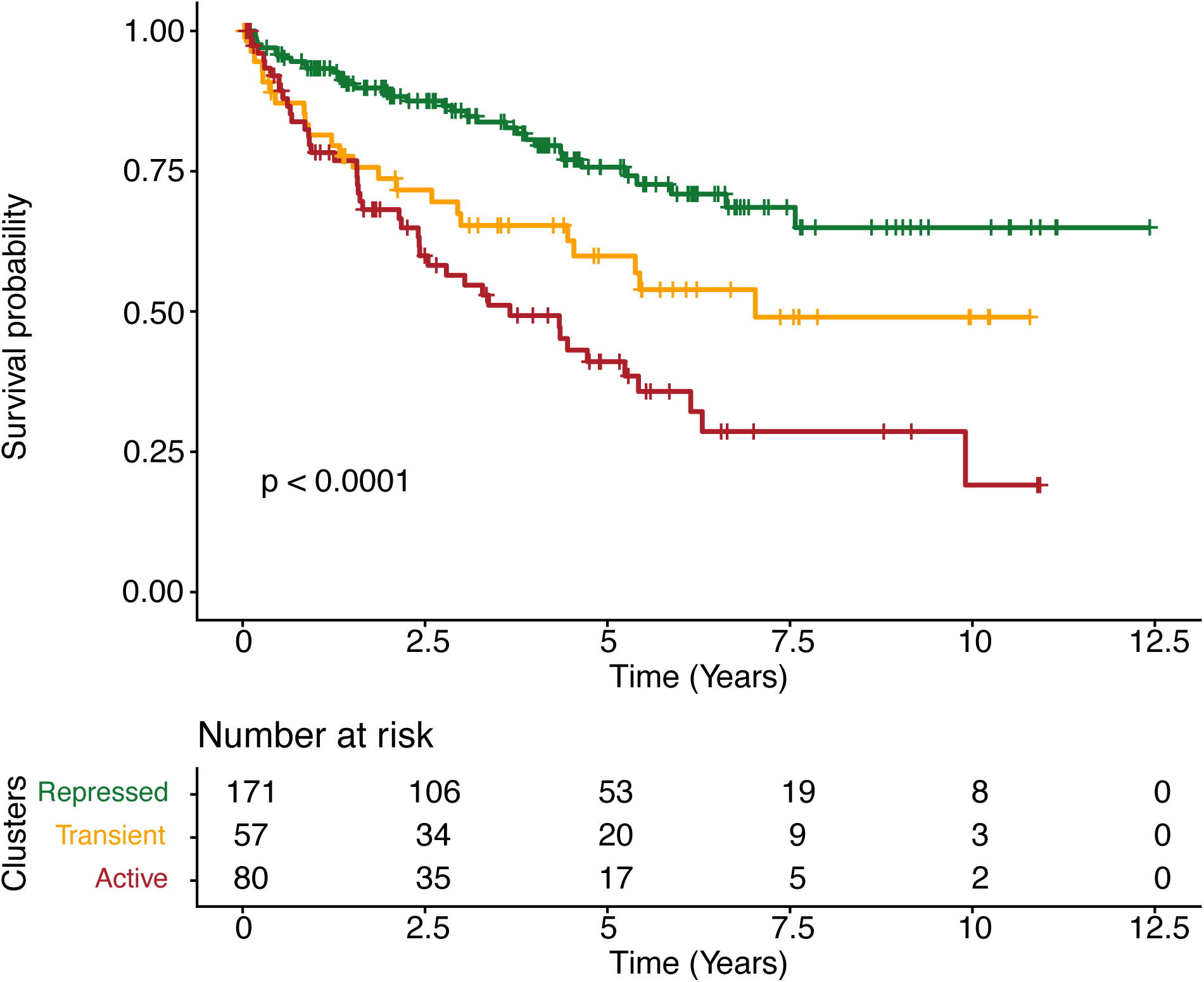
Active RE Methylation Subtype Predicts Poorer Survival Independent of Clinical and TME Composition. Kaplan–Meier survival curves for RE-defined tumor subgroups (Repressed, Transient, Active). The three subgroups demonstrate a graded survival pattern, with the Repressed group showing the most favorable outcomes, followed by Transient and Active tumors (log-rank *p* < 0.0001). Median overall survival was 3.7 years for the Active group (95% CI: 2.4–6.1) and 7.0 years for the Transient group (95% CI: 4.5–NA). Median survival was not reached for the Repressed group.

We next quantified these associations using multivariable Cox regression with the Repressed subtype as the reference (Table 3). After adjustment for age, sex, and AJCC stage (Model 1a), the Active subtype was associated with a significantly increased risk of all-cause mortality (HR = 1.85, 95% CI: 1.15–2.98, p = 0.011). A comparable effect was observed when AJCC stage was replaced by AUA risk group (Model 1b; HR = 1.81, 95% CI: 1.09–2.97, p = 0.011). To determine whether this association reflected differences in tumor microenvironment composition, we further adjusted for HiTIMED-derived immune and angiogenic components. Inclusion of these features did not significantly change the risk estimates for the Active subtype, either in the AJCC-adjusted model (Model 2a; HR =1.94, 95% CI: 1.21–3.12, p=0.006) or the AUA-adjusted model (Model 2b; HR = 1.97, 95% CI: 1.17-3.17, p= 0.0095). Across all model specifications, hazard ratios for the Active subtype remained consistently greater than one with confidence intervals that did not cross unity. In contrast, the Transient subtype showed intermediate risk estimates that were attenuated toward the null after adjustment and did not show a consistent independent association with survival.

**Table 3.** Summary of hazard ratios, 95% confidence intervals, and p values for the Active subtype across multivariable Cox models.

| Models | Covariates | HR | 95% CI | P value |
| --- | --- | --- | --- | --- |
| Model 1a | Age, Sex, Stage | 1.85 | 1.15-2.98 | 0.0114 |
| Model 1b | Model 1a + TME | 1.81 | 1.10-2.97 | 0.020 |
| Model 2a | Age, Sex, AUA Risk | 1.94 | 1.21-3.12 | 0.0060 |
| Model 2b | Model 2a +TME | 1.93 | 1.17-3.17 | 0.0095 |

Hazard ratio estimates remain consistent with and without adjustment for tumor microenvironment features, supporting an independent association between the Active RE methylation subtype and worse overall survival.

### 3.4. Differential methylation in the Active RE subtype converges on immune, cell-cycle, and metabolic genes

To investigate the gene-proximal consequences of RE demethylation, we performed an epigenome-wide association analysis comparing REMP-predicted RE CpGs between the Active and Repressed subtypes. Using |log2FC|>1 and FDR<0.05, we identified 25 differentially methylated positions (DMPs) (Figure 3). 22 CpGs were hypomethylated, and 3 (*LINC02C13, RAB11FIP3, RIMKLB*) were hypermethylated in the Active subtype, consistent with the global demethylation observed at the subtype level. After removing duplicated positions, these DMPs mapped to a focused set of genes enriched for immune regulation *(PIK3CD, HLA-E, HSPA1L*), cell-cycle control (*RASSF1*), Epithelial to Mesenchymal Transition (*ELMO1, RAB11FIP3*), and metabolism (Figure 3, Table 4, Table S2).

**Figure 3.**
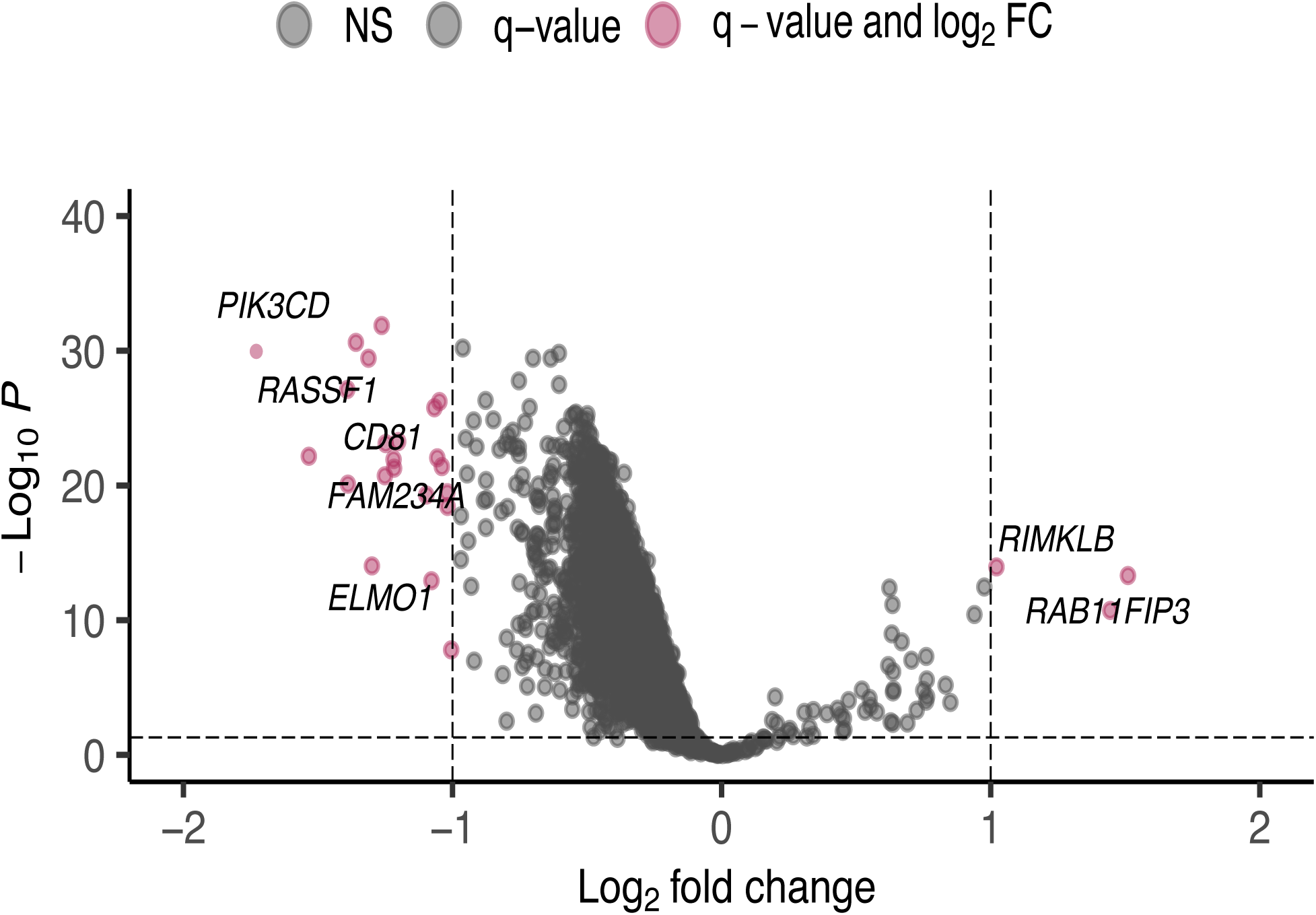
Differential Methylation Analysis Reveals Enrichment of Genes Related to Immune Response, Cell Cycle, Metabolic Genes in the Active RE Subset. Volcano plot showing differentially methylated RE-associated CpGs between the Active and Repressed subtypes (|log₂FC| > 1 and FDR < 0.05). Twenty-five positions were significantly altered, including 22 hypomethylated and 3 hypermethylated loci. Highlighted DMPs (pink) map to genes involved in immune activation (PIK3CD, CD81, HLA-E), cell-cycle regulation (RASSF1), EMT (ELMO1, RAB11FIP3), and metabolic pathways (SUCLG2-DT, HCG17, RPS18).

**Table 4.**
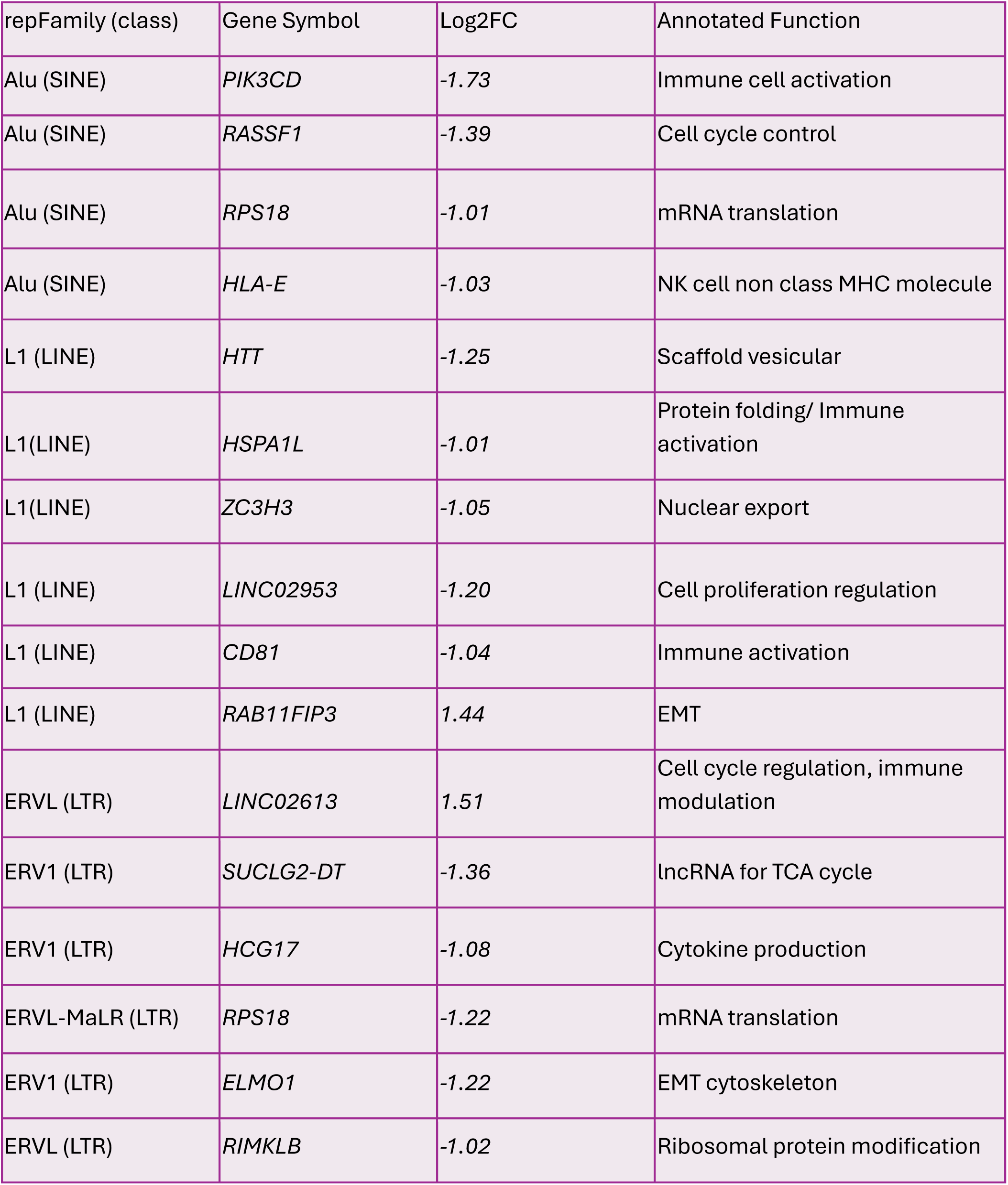
Top RE-linked CpGs, RE class/family, associated protein-coding or non-coding genes, and functional annotations.

### 3.5. RE hypomethylation is associated with transcriptional regulation of HIF2A expression and increased Immune response

We next examined whether RE methylation subtypes differed in hypoxia-associated gene expression and immune regulatory features, including interferon signaling. Across the three RE-defined groups, *EPAS1* (HIF2a) expression was significantly lower in the Active subtype compared with both the Repressed and Transient subgroups (both *p* < 0.001; Figure 4A). In contrast, *HIF1A* expression did not decrease in the Active group, with levels similar to or slightly higher than those in the other subtypes (Figure S3). Analysis of tumor microenvironment features revealed significantly higher Treg and CD8T cell abundance derived from HiTIMED in the Active subgroup (*p* < 0.0001 vs. Repressed/Transient; Figure 4B-C). Similarly, PD-1 (*PDCD1*) mRNA expression and T cell exhaustion signatures were also elevated in the Active subtype compared with both the Repressed and Transient groups (*p* < 0.001) for each comparison; Figure 4D-F). In contrast, RE methylation subtypes were not associated with the alteration status of common ccRCC mutational drivers *VHL, BAP1, PBRM1, and SETD2* (p = 0.062). (Figure S4).

**Figure 4.**
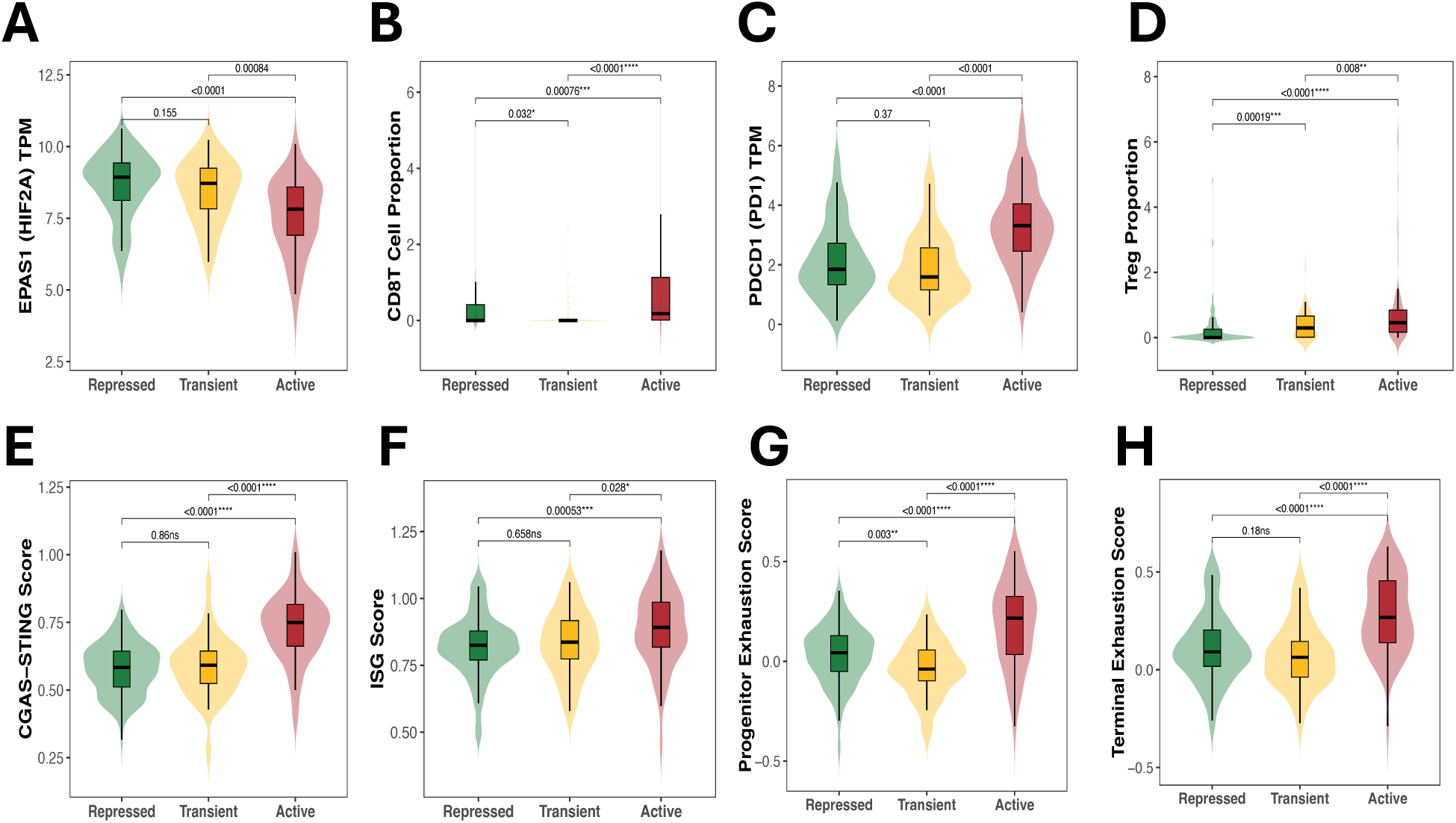
RE hypomethylation is associated with transcriptional reduction of HIF2A expression and increased Immune response. A *EPAS1* (HIF2A) mRNA expression across RE-defined subgroups. Active tumors show significantly lower *EPAS1* levels compared with both Repressed and Transient tumors. B CD8⁺ T cell i & Regulatory T cell (Treg) infiltration estimated from HiTIMED D Terminal T cell exhaustion scores compared among RE defined Methylation states. E *PDCD1* (PD-1) mRNA expression across subgroups. F cGAS–STING activation scores. G Interferon-stimulated gene (ISG) scores, as a proxy for enhanced interferon activity. H Progenitor exhaustion scores.

To assess associations with innate immune sensing pathways, we compared composite cGAS– STING activation and Interferon Stimulated Genes signature scores across the three subtypes. The Active subtype displayed significantly higher cGAS-STING ssGSEA scores than either of the Repressed and Transient groups (p < 0.0001 for each comparison), with a similar pattern for ISG scores (p < 0.05; Figure 4G-H).

### 3.6. External Validation of RE methylation subtypes in the CPTAC-ccRCC cohort reveals conserved cluster structure but altered global methylation ordering and context-dependent survival associations

To validate the RE methylation subtypes identified in TCGA-KIRC, we restricted the CPTAC methylation data to the 2,042 most variable RE-associated CpG positions that were shared between the cohorts. Using these common loci, NEMO integrative clustering (k = 3) revealed three distinct subgroups characterized by low, intermediate, and high global RE methylation levels, which we classified as Active (n = 78), Transient (n = 87), and Repressed (n = 78), respectively (Figure 5A). We next evaluated the prognostic relevance of the CPTAC RE subtypes using Cox proportional hazards models. Analyses were conducted in both the full cohort (n = 243; Repressed = 78, Transient = 87, Active = 78) and a complete-case subset with unique tumors and complete covariate information (n = 167; Repressed = 36, Transient = 46, Active = 85). To account for repeated tumor samples in CPTAC, we fit the model using cluster-robust standard errors in the full cohort (n = 243). In this model, the Transient subtype demonstrated independent prognostic significance relative to Repressed (HR = 4.8, 95% CI: 1.2–18.8, p = 0.03), whereas the Active subtype continued to trend toward worse outcomes (HR = 2.1, 95% CI: 0.8–5.3, p = 0.12) (Figure 5B). In the complete-case subset, the Active subtype showed a consistent trend toward inferior overall survival, mirroring TCGA results (Table S3).

**Figure 5.**
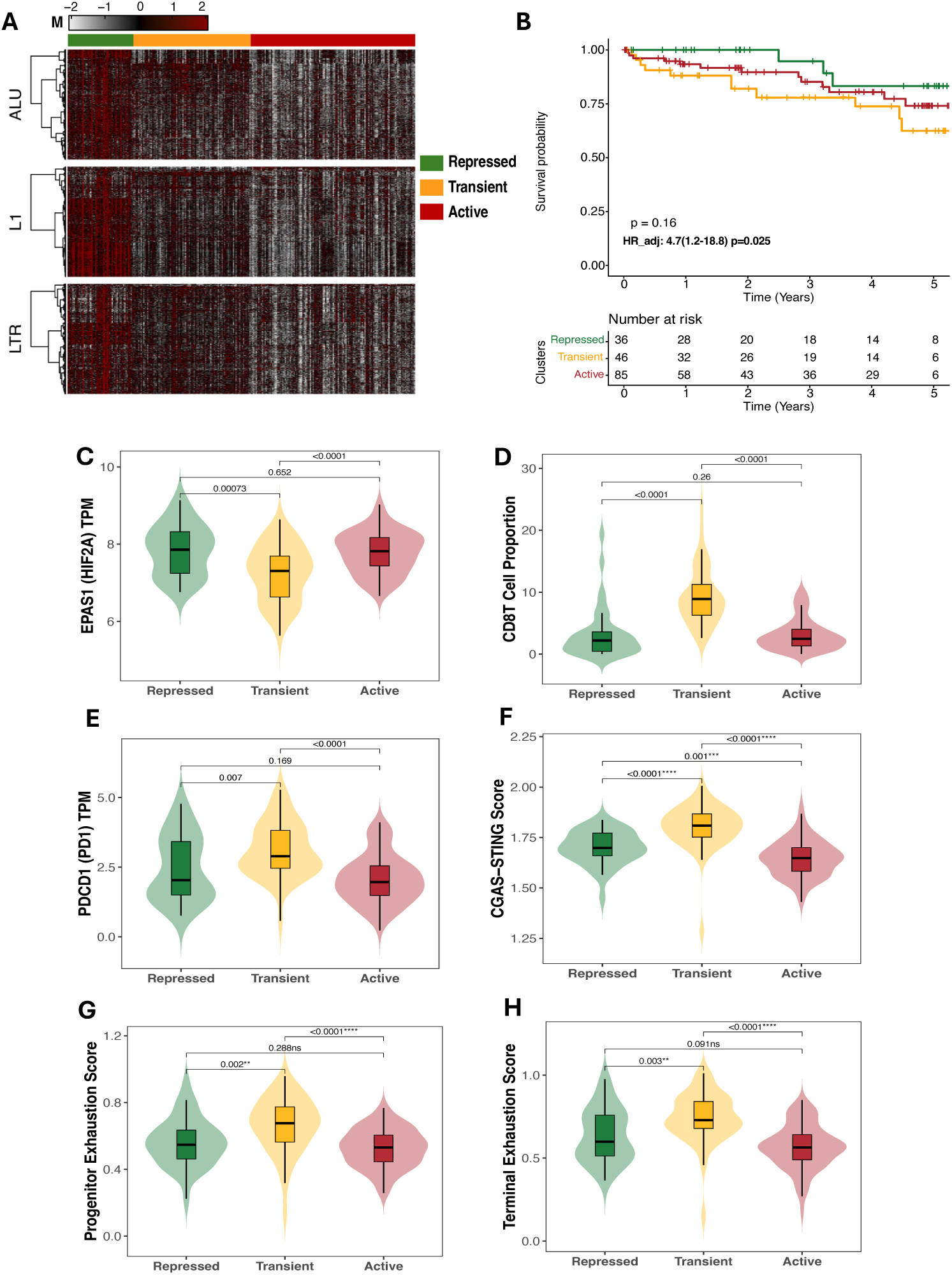
Validation of retroelement (RE) methylation subtypes in the CPTAC ccRCC cohort. **A** NEMO-based clustering (k=3) of REMP-predicted Alu, LINE-1 (L1), and LTR methylation in the CPTAC cohort identifies three RE-defined subtypes (Repressed, Transient, Active) with coordinated global methylation patterns consistent with those observed in TCGA. **B** Kaplan–Meier overall survival analysis shows separation of RE subtypes, with the Repressed group exhibiting the most favorable outcomes. Overall log-rank test significant (p = 0.16) and multivariable Cox regression adjusting for clinical covariates (HR_adj = 4.7, 95% CI: 1.2–18.8; p = 0.025) are shown. **C–H** Violin and box plots show molecular and immune features across RE subtypes, including *EPAS1* (HIF2A) mRNA expression, *PDCD1* (PD-1) mRNA expression, CD8⁺ T cell proportions, progenitor exhaustion scores, and terminal exhaustion scores. Box plots indicate median and interquartile range; violins show distribution density. Pairwise comparisons were performed using Wilcoxon rank-sum tests with multiple-testing adjustment, with adjusted p values shown.

Using CPTAC’s intrinsic RE methylation predictions (all variable loci rather than TCGA-shared loci), results were consistent. The Transient group remained independently prognostic for overall survival (HR = 9.06, 95% CI: 2.8–29) and progression-free survival (HR = 5.2, 95% CI: 1.9–13.5). In the complete-case subset, the Transient versus Repressed contrast also retained significance for overall survival (HR = 6.1, 95% CI: 1.4–26.2) (Figure 5B, Table S3). To identify common molecular alterations underlying the poorest-prognosis groups across datasets, we compared differential methylation results between high-risk subtypes in TCGA and CPTAC. Only one CpG position, ERV_0177340 (annotated to LINC01098), was shared and consistently hypomethylated (Table S2). We next evaluated whether transcriptomic correlates of RE hypomethylation observed in TCGA were recapitulated in CPTAC. In this cohort, the high-risk subtype (Transient) exhibited significantly reduced *EPAS1* (HIF2A) expression, elevated PD-1 expression, increased CD8 T-cell infiltration, and the highest cGAS–STING activation score relative to the low-risk Repressed group (Figure 5C-H). In contrast, differences between the Active and Repressed subtypes were not statistically significant for these features.

### 3.7. Internal validation in the DCC cohort confirms an Active RE methylation subtype with adverse prognosis and immune-exhausted tumor phenotype

After filtering, 509,008 CpGs for DCC dataset were used for RE methylation prediction and aggregation. A total of 63,625 RE positions were retained after aggregation, including 48,733 Alu, 6,477 L1, and 8,415 LTR positions. In addition, missing values, representing 2.9% of total beta values, were imputed.

To determine whether the RE methylation subtype structure observed in TCGA-KIRC and CPTAC-ccRCC could be reproduced in an institutional cohort, we applied NEMO integrative clustering to the Dartmouth Cancer Center (DCC) ccRCC cohort using REMP-predicted Alu, L1, and LTR methylation profiles. In contrast to the three-subtype structure identified in TCGA and CPTAC, clustering in DCC supported a two-group solution, resolving tumors into an Active and a Repressed RE methylation subtype (Figure 6A, Figure S5). The Active subgroup (n=45) showed lower global RE methylation across integrated RE classes, consistent with a derepressed retroelement methylation state, whereas the Repressed subgroup (n=71) retained comparatively higher RE methylation.

**Figure 6.**
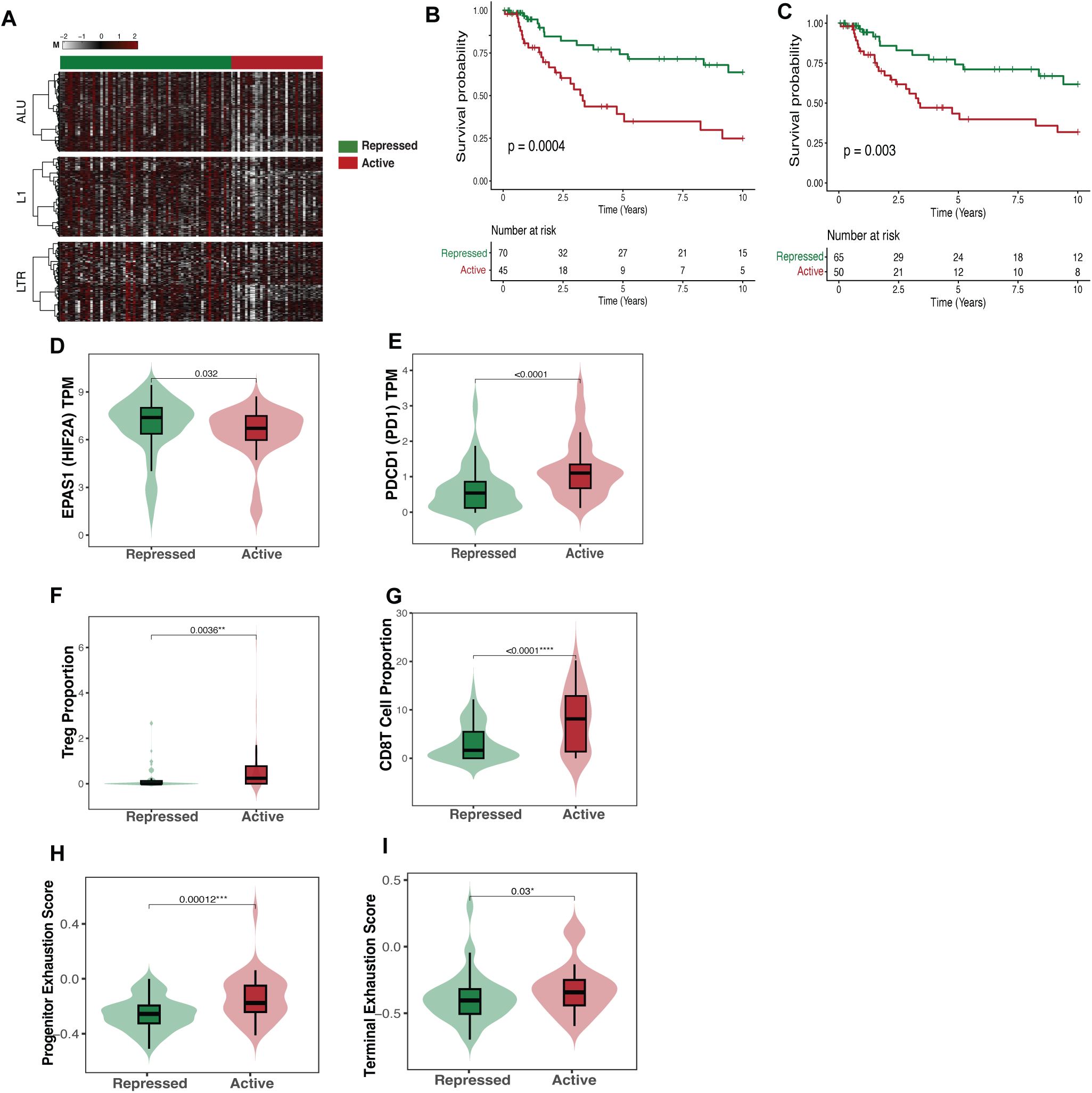
Independent validation and replication of retroelement (RE) methylation subtypes in the DCC ccRCC cohort. **A** NEMO-based clustering (k = 2) of REMP-predicted Alu, LINE-1 (L1), and LTR methylation in the Dartmouth Cancer Center (DCC) cohort identifies two RE-defined subtypes, Repressed and Active, with coordinated global methylation patterns across RE classes. **B** Kaplan, Meier overall survival analysis based on cohort-intrinsic DCC. & Kaplan, Meier overall survival analysis after restricting DCC clustering to RE-associated CpG loci shared with TCGA. **D-K** Violin and box plots show molecular and immune features across DCC RE subtypes, including **D** EPAS1 (HIF2A) mRNA expression, **E** PDCD1 (PD-1) mRNA expression, **F** Treg proportions, **G** CD8+ T cell proportions, **H** progenitor exhaustion scores, **I** terminal exhaustion scores.

Survival analysis demonstrated that the DCC Active subtype was associated with lower overall survival relative to the Repressed subtype (log-rank p= 0.0004; Figure 6B). However, this association was attenuated after adjustment for clinical and tumor microenvironment covariates, including age, sex, stage, and immune composition (HR = 1.07, 95% CI: 0.3 - 3.1, p = 0.9). To further harmonize the internal validation analysis with the TCGA discovery framework, we repeated clustering after restricting the DCC cohort to RE-associated CpG sites shared with TCGA. In this common-loci analysis, the Active subgroup showed a trend toward worse survival compared with the Repressed subgroup. Kaplan Meier analysis demonstrated significant unadjusted survival differences between the Active and Repressed subgroups (log-rank p = 0.003), but after multivariable adjustment the association was attenuated (HR = 1.09, 95% CI: 0.4 to 2.9, p = 0.8; Figure 6C)

We next evaluated whether molecular features associated with RE hypomethylation in TCGA were preserved in the DCC cohort. Consistent with the discovery analysis, *EPAS1* (HIF2A) expression was lower in the Active subgroup compared with the Repressed subgroup (Figure 6D). *HIF1A* expression showed a borderline association in the opposite direction, with higher expression in the Repressed subgroup (Figure S6).

We profiled immune signatures to explore the biological relevance of the DCC Active subtype. Active tumors showed increased CD8 T-cell and Treg infiltration compared with Repressed tumors (Figure 6E, F), accompanied by higher *PDCD1* expression (Figure 6G). More detailed immune-state analysis showed that both terminal exhaustion and progenitor exhaustion signatures were elevated in the Active subgroup (Figure 6H, I).

cGAS-STING scores did not differ significantly between Active and Repressed tumors. ISG scores were also not elevated in the Active subgroup but was higher in the Repressed subgroup (p = 0.024; Figure S6).

The DCC cohort reproduced the adverse prognostic association, reduced *EPAS1* expression, elevated *PDCD1* expression, and immune-exhausted phenotype of RE-hypomethylated tumors. Finally, differential methylation analysis comparing REMP-predicted RE CpGs between the Active and Repressed DCC subtypes identified 47 differentially methylated positions at |log2FC| > 1 and FDR < 0.05 (Table S2). Of these, 25 CpGs were hypomethylated in the Active subgroup, with top positions annotated to *VOPP1, SPINK2, and PIK3CD*, whereas 22 CpGs were hypermethylated with top positions annotated to *DEAF1, HYAL1, and DYRK1A*. Notably, *PIK3CD* was also hypomethylated in TCGA, whereas *LINC02C13* was among the hypermethylated loci identified in the TCGA analysis.

## Discussion

In this study, we integrated genome-wide DNA methylation predictions for Alu, LINE-1, and LTR retroelements to define epigenetic subtypes of clear cell renal cell carcinoma and evaluated their molecular, immunologic, and clinical relevance across a discovery cohort, an external validation cohort, and an independent replication cohort. Three reproducible RE methylation subtypes emerged in TCGA-KIRC, designated Repressed, Transient, and Active, with the Active subtype showing the most extensive hypomethylation across RE classes and the clearest adverse clinical profile. Patients in the Active group had significantly worse survival than those in the Repressed group, and this association remained evident after adjustment for clinicopathologic and tumor microenvironment variables. To reduce the influence of competing non-cancer mortality in the longer-followed cohorts, survival analyses in TCGA and DCC were censored at 10 years. Together, these findings indicate that RE methylation captures a clinically meaningful dimension of tumor aggressiveness beyond established prognostic models. Prior work linking ERV hypomethylation to genomic instability and poor outcome in ccRCC is consistent with this interpretation, and our results extend that concept to a genome-wide RE methylation framework spanning multiple retroelement classes^55^.

The biological features of the Active subtype further support the relevance of this RE methylation axis. In TCGA, RE hypomethylation was associated with reduced *EPAS1* expression, increased CD8 T-cell and Treg infiltration, elevated *PDCD1* expression, and higher progenitor and terminal exhaustion scores, together with increased cGAS-STING and interferon-stimulated gene activity. This pattern is consistent with an inflamed but ineffective tumor immune microenvironment, in which antiviral-like signaling coexists with immune suppression and T-cell dysfunction. Prior studies have similarly shown that endogenous retroviral activity can promote immune activation while also contributing to checkpoint expression and dysfunctional antitumor immunity in ccRCC^6,6,23^. The coordinated activation of cGAS-STING, interferon-related programs, and checkpoint pathways therefore provides orthogonal support for the biological significance of RE hypomethylation, even though retroelement transcription was not directly measured^13,25,52–53^.

The association between the Active subtype and reduced EPAS1 mRNA expression is also notable. Although HIF2α is a central driver of ccRCC biology, its prognostic significance appears to depend on whether mRNA or protein is being measured, as well as on cellular context and protein localization. Our finding that lower *EPAS1* transcript abundance marks the poorest-prognosis RE hypomethylated tumors is therefore consistent with prior reports linking reduced *EPAS1* mRNA to aggressive disease behavior in ccRCC.^56–58^

External validation in CPTAC largely supported the robustness of this framework, while also highlighting expected cohort-specific variation. The overall survival pattern remained directionally consistent, with the more repressed state showing the most favorable outcomes, although in CPTAC the Transient subtype emerged as the clearest high-risk group. Given the lower number of events, shorter follow-up, and very low event rate in the Repressed reference group, this difference likely reflects cohort structure rather than instability of the underlying RE methylation signal. Importantly, key molecular correlates of the high-risk state were preserved in CPTAC, including reduced *EPAS1* expression, elevated PD-1 expression, increased CD8 infiltration, and heightened cGAS-STING activity.

The independent validation and replication analysis in the DCC cohort further strengthened the overall framework. Unlike TCGA and CPTAC, where integrative clustering resolved three subtypes, the DCC cohort supported a two-group solution separating tumors into Active and Repressed states, likely reflecting limited power to resolve a distinct intermediate state. Even within this simplified structure, the key biological features of the Active state were recapitulated. Active tumors showed poorer unadjusted survival, reduced *EPAS1* expression, increased *PDCD1* expression, higher CD8 T-cell and Treg infiltration, and elevated progenitor and terminal exhaustion signatures, supporting the reproducibility of the immune-exhausted phenotype across cohorts.

At the same time, the DCC results underscore that replication should be interpreted at the level of biological pattern rather than exact numerical equivalence. Although both TCGA and DCC used TPM-based expression data, absolute expression ranges for *EPAS1, PDCD1*, and derived signature scores differed between cohorts. This is expected because TPM values are shaped by cohort-specific expression distributions, RNA-seq processing workflows, sample composition, and tumor purity, while ssGSEA-like scores are relative to the internal expression background of each dataset. Accordingly, concordance is best assessed through directionality and subtype separation within each cohort rather than by matching absolute score magnitudes across studies.

One notable difference in DCC concerned interferon-related signaling. In TCGA, the Active subtype showed higher cGAS-STING and ISG activity, consistent with a viral mimicry-like state associated with RE derepression. In DCC, however, cGAS-STING scores were not significantly different between groups, and ISG scores were higher in the Repressed subgroup. This suggests that while RE hypomethylation reproducibly marks a poor-prognosis, immune-exhausted phenotype, its coupling to downstream innate immune activation may be more context dependent. Smaller sample size, reduced power, the two-group subtype structure, and cohort-specific technical or clinical differences may all contribute. Biologically, these findings raise the possibility that chronic T-cell dysfunction is a more stable correlate of RE hypomethylation than interferon activation itself.

Differential methylation analysis further suggested that the Active RE state is accompanied not only by broad retroelement hypomethylation, but also by more focal epigenetic changes at genes involved in tumor stress adaptation, matrix remodeling, and intracellular signaling, all of which are relevant to tumor progression and microenvironmental interaction (Table S2). Consistent with this, comparison of the high-risk groups in TCGA and CPTAC identified a shared hypomethylated locus, *ERV_0177340* annotated to *LINC010S8*, supporting the idea that focal RE-associated methylation changes accompany the broader subtype structure. In DCC, the recurrence of *PIK3CD* and *LINC02C13*, also identified in TCGA, suggests that despite broader differences in global RE methylation patterns, a small subset of gene-proximal loci may mark conserved features of the high-risk epigenetic state. This is biologically notable because *PIK3CD* is linked to immune signaling, whereas *LINC02C13* may reflect noncoding regulatory remodeling, together reinforcing the idea that RE hypomethylation co-occurs with focal epigenetic changes at loci relevant to tumor-immune interactions and aggressive disease behavior.

This study has several limitations. Retroelement methylation was inferred using the REMP framework rather than measured directly at each locus. Although validated, REMP provides genome-wide estimates from array data and may miss nuanced regulatory effects. Survival analyses in CPTAC were limited by fewer events, a younger cohort with shorter follow-up, and low event rates in the Repressed reference group, reducing power despite consistent trends. In DCC, the smaller sample size and two-group solution may likewise have limited power to resolve an intermediate subtype and detect subtler downstream transcriptional differences, particularly for interferon-related signatures. Cross-cohort differences in RNA-seq processing, sample composition, and clinical context also likely contributed to differences in absolute expression and signature score scales.

Overall, our findings define a reproducible RE-centered epigenetic framework for ccRCC with clear biological and clinical relevance. Across discovery, external validation, and independent institutional replication cohorts, tumors with more active RE methylation states consistently showed features of aggressive disease, immune dysfunction, and poorer outcome. These results support RE methylation profiles as a meaningful dimension of ccRCC heterogeneity and a potential complement to existing molecular classification strategies.

## Conclusions

We identified retroelement methylation subtypes in ccRCC that capture a biologically meaningful axis of tumor variation linked to immune state and clinical outcome. Across cohorts, tumors with more derepressed RE methylation profiles consistently showed features of aggressive disease, including reduced *EPAS1* expression, heightened immune dysfunction, and poorer survival in the discovery setting. External validation and independent replication supported the broader biological relevance of this framework, even where the strength of prognostic associations varied after adjustment. These findings position RE methylation as an informative layer of ccRCC heterogeneity that complements established clinicopathologic and molecular features. More broadly, they suggest that retroelement dysregulation may help connect epigenomic instability to immune remodeling and adverse tumor behavior in ccRCC.

## List of abbreviations

AJCC: American Joint Committee on Cancer
ANOVA: Analysis of Variance
AUA: American Urological Association
ccRCC: Clear Cell Renal Cell Carcinoma
CNA: Copy Number Alteration
COCA: Cluster-of-Clusters Analysis
CPI: Cluster Prediction Index
CPTAC: Clinical Proteomic Tumor Analysis Consortium
NAm: DNA Methylation
EPAS1: Endothelial PAS Domain Protein 1
FDR: False Discovery Rate
GSVA: Gene Set Variation Analysis
HERV: Human Endogenous Retrovirus
HIF: Hypoxia-Inducible Factor
HiTAiC: Hierarchical Tumor Artificial Intelligence Classifier
HiTIMED: Hierarchical Tumor Immune Microenvironment Epigenetic Deconvolution
HR: Hazard Ratio
IntNMF: Integrated Non-Negative Matrix Factorization
ISG: Interferon-Stimulated Gene
KNN: k-Nearest Neighbors
L1: LINE-1
LINE: Long Interspersed Nuclear Element
LTR: Long Terminal Repeat
MAD: Median Absolute Deviation
MOVICS: Multi-Omics Integration and Clustering System
NEMO: Neighborhood-based Multi-Omics Clustering
PD-1: Programmed Cell Death Protein 1
PDCD1: Programmed Cell Death 1 Gene
RE: Retroelements
REMP: Repetitive Element Methylation Prediction
SNF: Similarity Network Fusion
SNP: Single Nucleotide Polymorphism
ssGSEA: Single-Sample Gene Set Enrichment Analysis
TCGA-KIRC: The Cancer Genome Atlas Kidney Renal Clear Cell Carcinoma
TME: Tumor Microenvironment
TPM: Transcripts Per Million
VHL: von Hippel–Lindau

## Declarations Authors Affiliation

Chinaza F. Nnam Department of Cancer Biology, Molecular and Cellular Biology PhD Program, Dartmouth College

Erick A. Mboya, Department of Epidemiology, Quantitative Biomedical Sciences PhD Program, Dartmouth College

Patricia A. Pioli, Associate Professor of Microbiology and Immunology. Dartmouth College

Li Song, Assistant Professor of Biomedical Data Science, Assistant Professor of Microbiology and Immunology. Dartmouth College

Brock C. Christensen, Professor of Epidemiology, Geisel School of Medicine at Dartmouth. Professor of Molecular and Systems Biology. Dartmouth College

Lucas A. Salas, Assistant Professor of Epidemiology, Geisel School of Medicine at Dartmouth. Assistant Professor of Cancer Biology. Dartmouth College

John D. Seigne: Department of Surgery, Section of Urology, Geisel School of Medicine, Dartmouth College, Lebanon, NH 03766, USA.

Jason R. Pettus Department of Pathology and Laboratory Medicine, Dartmouth Hitchcock Medical Center, Lebanon, NH, USA. Geisel School of Medicine at Dartmouth, Hanover, NH, USA.

Bing Ren Department of Pathology and Laboratory Medicine, Dartmouth-Hitchcock Medical Center, Lebanon, New Hampshire.

Marc Ernstoff. Director, Experimental Cell Therapy Program. Professor of Medicine Geisel School of Medicine at Dartmouth. Dartmouth-Hitchcock Medical Center Lebanon, New Hampshire.

## Ethics approval

Not applicable

## Consent for publication

Not applicable

## Availability of data and materials

The datasets analyzed during the current study are publicly available from the Genomic Data Commons (GDC) Data Portal under the TCGA Kidney Renal Clear Cell Carcinoma (TCGA-KIRC) project (https://portal.gdc.cancer.gov project id: TCGA-KIRC). CPTAC clear cell renal cell carcinoma (CPTAC-3 ccRCC) data are available through the Clinical Proteomic Tumor Analysis Consortium via the GDC Data Portal and the CPTAC Data Portal (https://portal.gdc.cancer.gov project id: CPTAC-3), subject to applicable data access policies. The analysis scripts used in this study are available in https://github.com/SalasLab/Retroelements_Methylation_Analysis_in_ccRCC)

## Authors’ contributions

Conceptualization: CFN, LAS; Methodology: CFN, LAS; Formal Analysis: CFN, EAM; Investigation: CFN;Visualization: CFN; Writing – Original Draft: CFN; Writing – Review & Editing: CFN, EAM, PAP, ME, JDS, JRP, BR, LS, BCC, LAS; Supervision: LAS; Funding Acquisition: LAS.

## Additional files

**Additional file 1**: Table S1A: cGAS–STING gene set. Table S1B: Interferon-stimulated gene set. Table S1C: Terminal exhaustion gene set. Table S1D: Progenitor exhaustion gene set.

Table S2A: Differentially methylated positions between Active and Repressed RE groups in the TCGA cohort. Table S2B: Common differentially methylated positions identified in both TCGA-KIRC, CPTAC-ccRCC and DCC analyses.

Table S3A: Hazard ratios and 95% confidence intervals based on clustering of common TCGA and CPTAC-derived beta values. Table S3B: Hazard ratios and 95% confidence intervals based on clustering of CPTAC-only derived beta values. Table S3C: Hazard ratios and 95% confidence intervals for RE-defined subgroups in DCC cohort.

**Additional file 2**: Figure S1: Benchmarking of integrative clustering approaches for retroelement methylation data. Figure S2: Comparison of integrative clustering algorithms applied to retroelement methylation profiles. Figure S3: *HIF1A* mRNA expression across RE-defined tumor subgroups in discovery and validation cohorts. Figure S4: Associations between retroelement methylation subtypes and key molecular drivers in clear cell renal cell carcinoma. Figure S5: RE Methylation-Based Clustering in the DCC Cohort. Figure S6: *HIF1A* (HIF-1α) mRNA expression, ISG score and cGAS-STING scores across RE-defined tumor subgroups in DCC cohort.

