## Supplementary Figures (Fig s1-s6) for "Retroelement Hypomethylation Links Hypoxia Signaling, Immune Phenotypes, and Survival in Clear Cell Renal Cell Carcinoma"

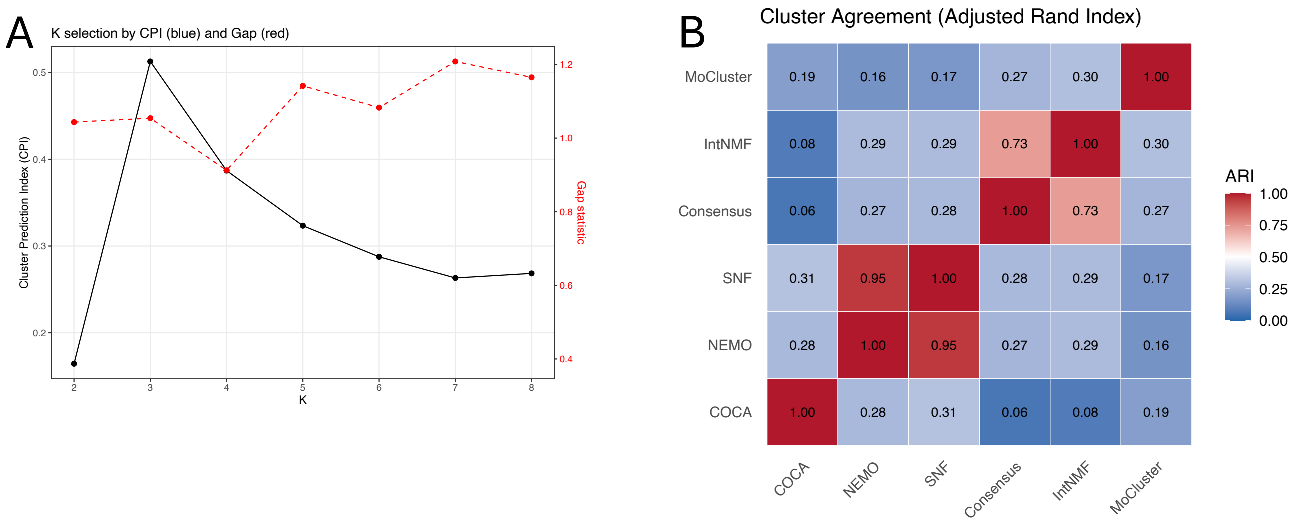


**Figure S1. Benchmarking of integrative clustering approaches for retroelement (RE) methylation data.**

**(A)** Determination of the optimal number of clusters (k) using the gap statistic and cluster prediction index (CPI) across integrated Alu, LINE-1, and LTR methylation profiles. CPI metric supported k = 3 as the most stable and informative solution, balancing cluster separation and robustness.

**(B)** Pairwise cluster agreement across six integrative clustering methods, quantified using the adjusted Rand index (ARI). Heatmap values indicate the degree of concordance between clustering solutions, with higher ARI reflecting stronger agreement. Network-based methods, particularly Neighborhood-based Multi-Omics clustering (NEMO) and Similarity Network Fusion (SNF), demonstrated the highest concordance and stability, whereas consensus- and matrix factorization–based approaches showed greater variability. These results motivated selection of NEMO for downstream subtype definition and clinical association analyses.

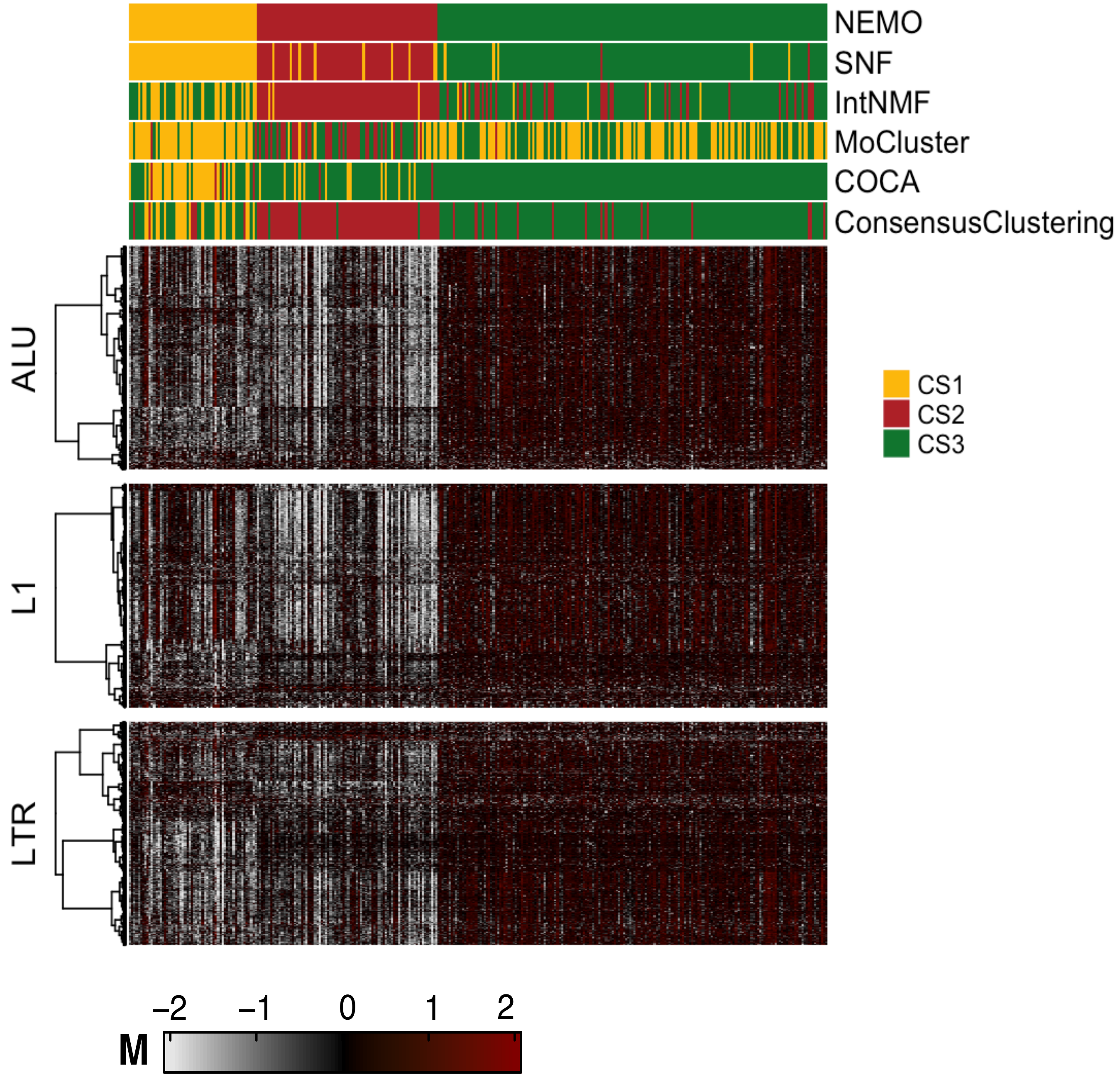


**Figure S2. Integrative clustering algorithms applied to retroelement (RE) methylation profiles.** Heatmaps display integrated REMP predicted DNA methylation levels across Alu, LINE 1, and LTR elements in the TCGA-KIRC cohort. Samples are annotated by cluster assignments (CS1-3) derived from six multi omics clustering frameworks, NEMO, SNF, IntNMF, MoCluster, COCA, and Consensus Clustering, each evaluated at k = 3. The NEMO Cluster (CS) assignment was later to Repressed (CS3), Transient (CS1), and Active (CS2). Color scale represents relative methylation levels (M values).

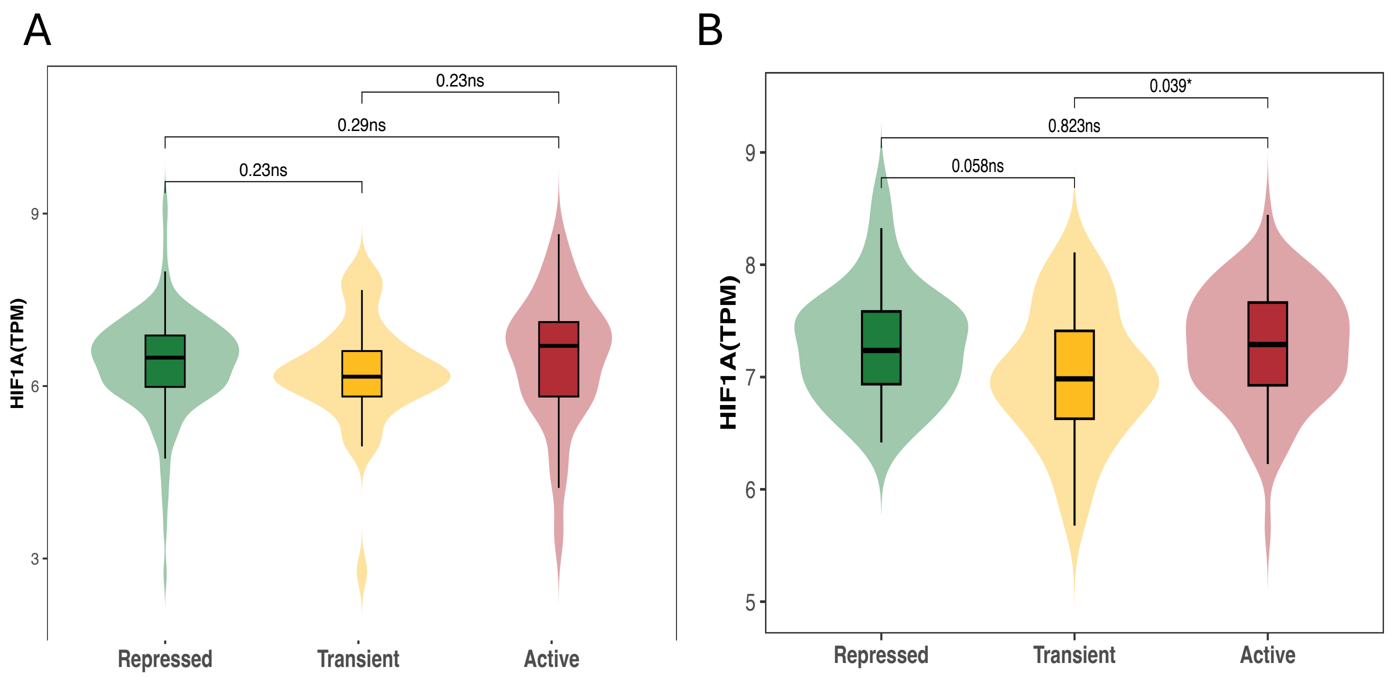


**Figure S3. HIF1A (HIF-1α) mRNA expression across RE-defined tumor subgroups in discovery and validation cohorts.**

Violin plots show HIF1A mRNA expression (TPM) across Repressed, Transient, and Active RE methylation subtypes in the TCGA discovery cohort (A) and the CPTAC validation cohort (B). In TCGA, HIF1A expression did not differ significantly between the low-risk Repressed and high-risk Active subgroups, with no significant pairwise comparisons observed. In CPTAC, HIF1A expression differed significantly across subgroups overall, driven primarily by differences involving the Transient subgroup, while the Active versus Repressed comparison remained non-significant. P values shown correspond to pairwise Wilcoxon rank-sum tests with multiple-testing adjustment, as indicated. These results suggest cohort-dependent variation in HIF1A expression that does not consistently distinguish low- and high-risk RE methylation states.

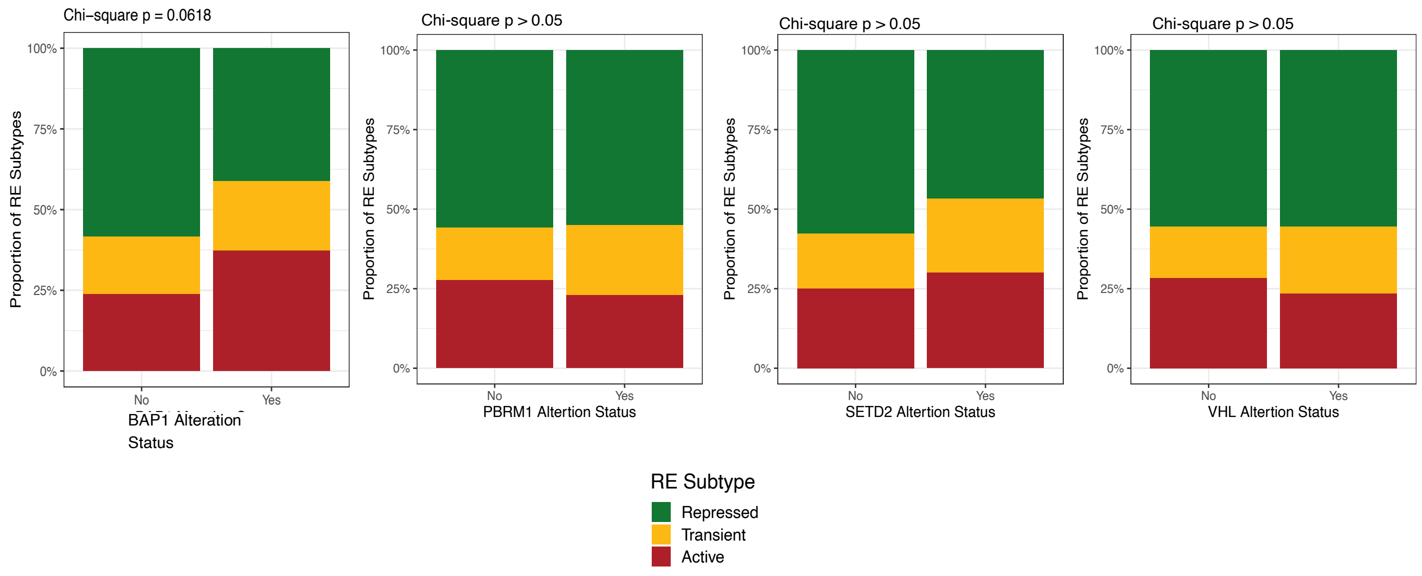


**Figure S4. Associations between retroelement methylation subtypes and key molecular drivers in clear cell renal cell carcinoma.**

Stacked bar plots showing the distribution of retroelement (RE) methylation subtypes (Repressed, Transient, and Active) according to alteration status of major ccRCC driver genes. Alteration status (Yes vs No) is shown for *BAP1*, *PBRM1*, *SETD2*, and *VHL*. Bars represent the proportion of tumors within each molecular driver assigned to each RE-defined subgroup. Statistical significance was assessed using chi-square tests. RE subtype composition trended towards statistical significance for differences in *BAP1* alteration status (chi-square p = 0.0618), while distributions across other driver genes are shown for comparison.

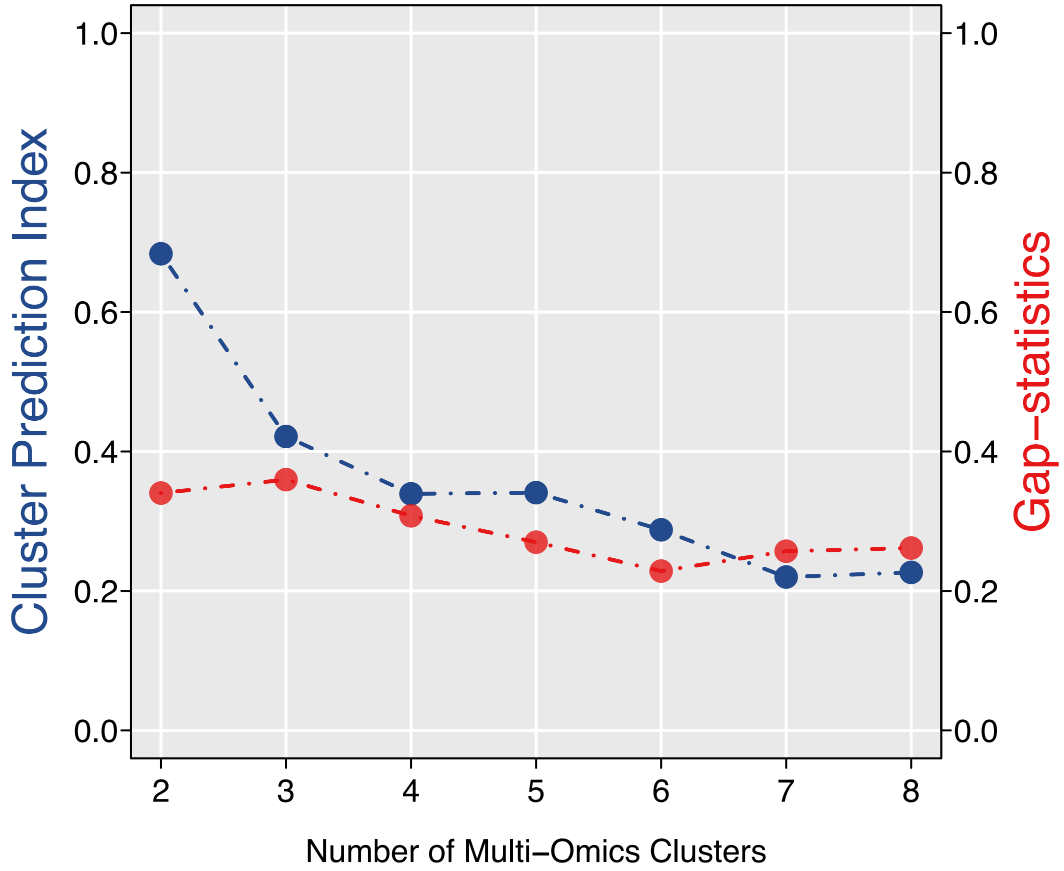


**Figure S5. RE Methylation-Based Clustering in DCC Cohort.**
CPI metric supported k = 2 as the most stable and informative solution, balancing cluster separation and robustness.

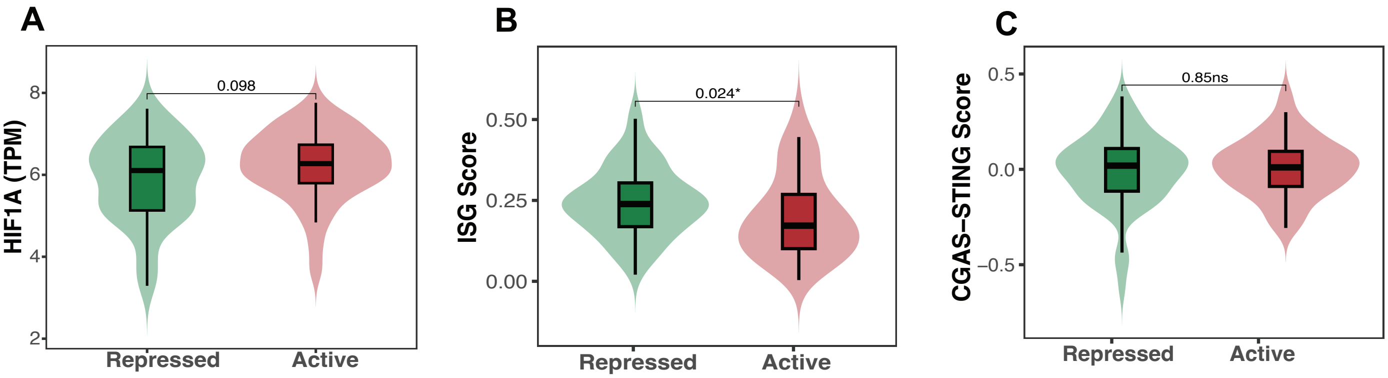


**Figure S6.** ***HIF1A*** **(HIF-1α) mRNA expression, ISG score and cGAS-STING scores across RE-defined tumor subgroups in DCC cohort.**

Violin plots show **HIF1A** mRNA expression (TPM) **(A)**, **interferon-stimulated gene (ISG) score (B)**, and **cGAS-STING score (C)** across the **Repressed** and **Active** retroelement (RE) methylation subgroups identified in the Dartmouth Cancer Center (DCC) cohort. HIF1A expression showed a nonsignificant trend across subgroups, whereas the ISG score was higher in the Repressed group. In contrast, cGAS-STING scores did not differ significantly between the Repressed and Active subgroups. P values shown correspond to Wilcoxon rank-sum tests.
